# Surface productivity gradients govern changes in abundance and physiological status of deep ocean prokaryotes across the tropical and subtropical Atlantic

**DOI:** 10.1101/2022.06.20.496789

**Authors:** Markel Gómez-Letona, Javier Arístegui, Nauzet Hernández-Hernández, María Pérez-Lorenzo, Xosé Antón Álvarez-Salgado, Eva Teira, Marta Sebastián

## Abstract

Prokaryotes represent a major fraction of marine biomass and play a key role in the global carbon cycle. We studied the vertical profiles (from surface down to the bathypelagic realm) of abundance, cytometric signatures, and activity of prokaryotic communities along a productivity gradient in the subtropical and tropical Atlantic to assess whether there is a vertical linkage between surface productivity regimes and deep ocean prokaryotic communities. We found that latitudinal changes in the vertical patterns of cytometric variables were coupled with surface productivity: higher prokaryotic abundances and viabilities, and smaller cell sizes were observed below highly productive surface waters, an effect reaching down to the bathypelagic layer. On the contrary, leucine uptake rates in deep waters showed no clear relationship with surface productivity. Changes in resource and energy allocation to growth vs. maintenance in hostile environments, cell-size-dependent metabolic requirements and variability in leucine to carbon conversion may all be part of the array of factors involved in controlling prokaryotic activity patterns that were measured. Our work adds to the recent findings that highlight the importance of vertical connectivity for prokaryotic communities in the dark ocean.

## Introduction

Prokaryotes represent a key component of marine ecosystems. At abundances typically ranging from thousands to millions of cells per millilitre, they conform a major fraction of the biomass of marine organisms (Whitman et al. 1998; Bar-On et al. 2018) and make use of a wide variety of energy and carbon sources (Moran 2015). The activity of heterotrophic prokaryotes plays a crucial role in global biogeochemical cycles and marine food webs. Consuming organic matter, they remineralise the chemical elements to inorganic forms, making them readily available for new cycles of primary production. They also channel organic matter into the marine food web by means of secondary production, i.e., the creation of heterotrophic biomass (Azam and Malfatti 2007), and generate an extremely diverse array of DOM molecules as a by-product of their metabolism (Dittmar et al. 2021). While prokaryotes are distributed along the entire water column, their abundance and activity are not constant. Epipelagic communities exhibit markedly higher abundances, biomass and production rates, lower percentages of cells with high nucleic acid content, and smaller cell sizes than those in meso- and bathypelagic waters (Arístegui et al. 2009). Nonetheless, given the vast volume of water encompassed by the dark ocean, prokaryotes in this realm are responsible for 75% and 50% of the ocean’s prokaryotic biomass and production, respectively (Arístegui et al. 2009), underlining the importance of considering the communities in the entire water column when studying carbon fluxes in the ocean.

In surface open-ocean waters the organic matter consumed by prokaryotes stems either from *in situ* primary production or from lateral advection from coastal adjacent regions (Santana-Falcón et al. 2020). In the dark ocean, however, while chemolithoautotrophy (Baltar et al. 2010b) or carbon excretion by diel vertical migrants (Steinberg et al. 2008) may significantly contribute as carbon sources, the vertical flux of particles that escape remineralisation in the photic layer is assumed to be the main source of organic carbon (Boyd et al. 2019). These particles have diverse origins: they may consist of phytoplankton cells (Guidi et al. 2009), zooplankton faecal pellets (Turner 2015) or polymer gel structures (Verdugo 2012). Not only they are a source of organic matter to deep ocean layers, but also act as vectors that vertically connect prokaryotic communities (Mestre et al. 2018; Ruiz-González et al. 2020). However, a link between surface productivity or particle flux and prokaryotic biomass or production in the dark ocean has not always been found (Arístegui et al. 2009 and references therein; Herndl and Reinthaler 2013), suggesting that differences might exist between oceanic provinces and/or times of the year.

In the present study, we analyse quantitative and qualitative cytometric signatures and heterotrophic activity estimates of prokaryotic communities along a section crossing the tropical and subtropical Atlantic Ocean, both in the epipelagic layer (0-200 m) and in the deep ocean, down to 3500 m. The studied region is complex, with a considerable number of currents (Brandt et al. 2008) and water masses (Pérez et al. 2001; Álvarez et al. 2014). Moreover, it presents a strong surface productivity gradient, comprising both oligotrophic waters and areas directly under the influence of the Northwest African upwelling system, where high export rates of sinking particles have been reported (Fischer et al. 2020). We determined the abundance and viability of prokaryotic communities, estimated their biomass, and measured leucine incorporation rates, to assess whether these variables were affected by changes in surface productivity, evaluating the degree to which deep prokaryotic communities were linked to epipelagic waters.

## Methods

### Study area

During the MAFIA cruise (*Migrants and Active Flux In the Atlantic ocean*, April 2015) seawater samples were collected at 13 stations along a section crossing a productivity gradient in the subtropical and tropical Atlantic (13°S –27°N, Fig. 1; Gómez-Letona et al., submitted). Sample collection was performed using a General Oceanics oceanographic rosette equipped with 24-L PVC Niskin bottles alongside a Seabird 911-plus CTD, a Seapoint Chlorophyll Fluorometer and a Seabird-43 Dissolved Oxygen Sensor. The chlorophyll-a (Chl-a) values provided by the fluorometer were based on the factory calibration.

**Fig. 1.**
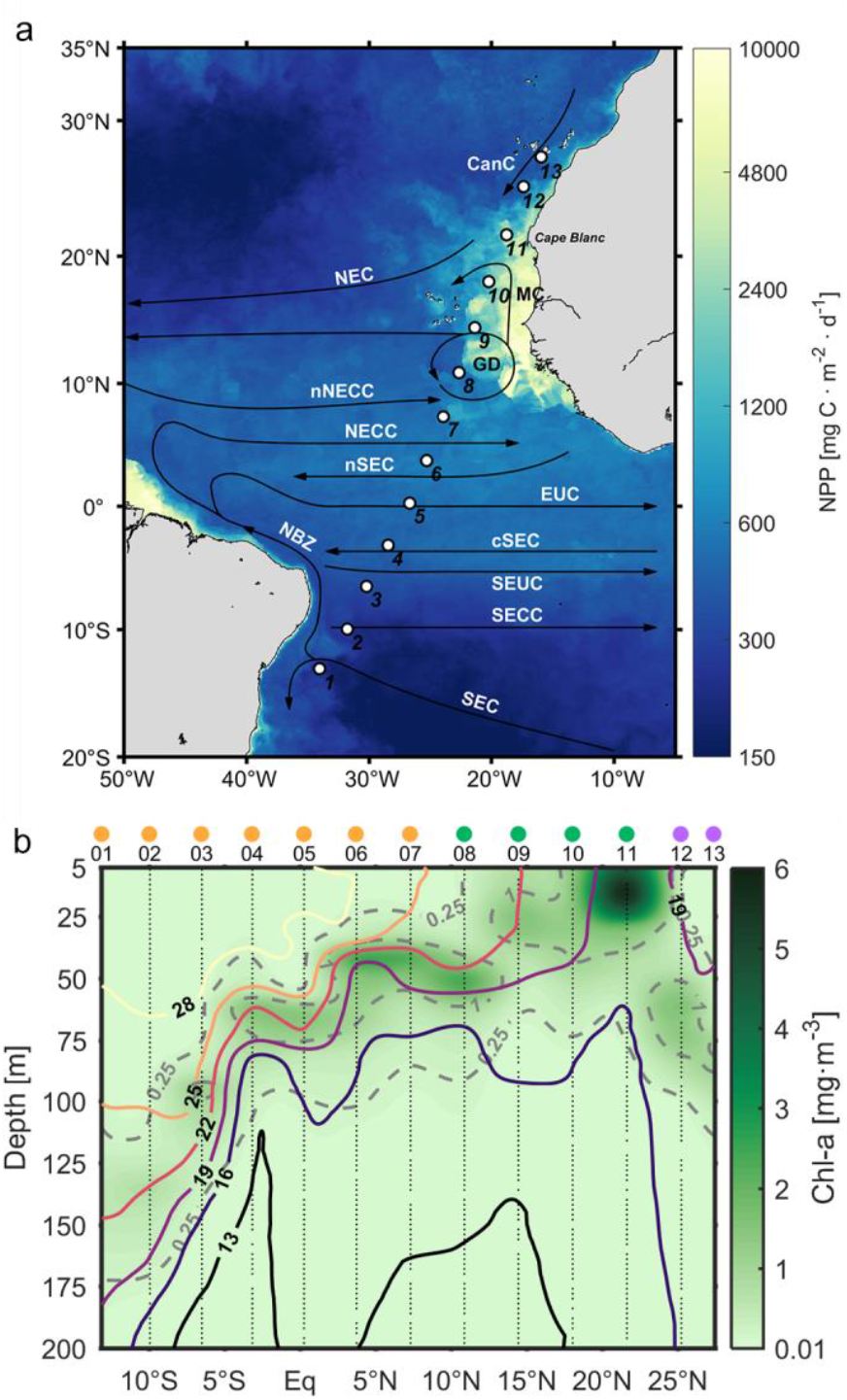
Surface productivity during the MAFIA cruise. (a) Sampling stations (1-13). The underlying colormap represents net primary production estimates for April 2015 (Eppley-VGPM model, MODIS dataset, Oregon State University, http://sites.science.oregonstate.edu/ocean.productivity/). Major ocean currents in the study region (based on Stramma and Schott, 1999 and Brandt et al., 2008) are also displayed: CanC, Canary Current; NEC, North Equatorial Current; MC, Mauritanian Current; GD, Guinea Dome; nNECC, northern North Equatorial Counter Current; NECC, North Equatorial Counter Current; nSEC, northern South Equatorial Current; EUC, Equatorial Under Current; NBZ, North Brazil Current; cSEC, central South Equatorial Current; SEUC, South Equatorial Under Current; SECC, South Equatorial Counter Current; SEC, South Equatorial Current. (b) Chl-a concentrations (derived from the CTD fluorometer) and potential temperature isotherms (in °C) in the epipelagic layer along the cruise section. Dashed isolines represent Chl-a concentrations. Dots on top of station numbers represent station groups: ‘South’ (light orange), Guinea Dome–Cape Blanc (‘GD-CB’, green), ‘North’ (violet).

### Prokaryotic cell abundance and viability

Seawater samples for measuring the abundance of prokaryotes were collected in all stations at 22 depths, from surface down to 3500 m (or bottom, when above this depth). Samples were collected into 1.2 mL cryovials, fixed with a 2% final concentration of formaldehyde, after keeping them 30 min at 4°C, and then stored frozen in liquid nitrogen. After 24 h they were analysed in a FACSCalibur (Becton-Dickinson) flow cytometer. Subsamples (400 μL) were stained with 4 μL of the fluorochrome SYBR Green I (Molecular Probes) diluted in dimethyl sulfoxide (1:10). Fluorescent beads (1 μm, Polysciences) were added for internal calibration (10^5^ mL^−1^). High and low nucleic acid content (HNA and LNA, respectively) prokaryotic cells were identified in green (FL1) vs red fluorescence (FL3) and FL1 vs side scatter (SSC) cytograms. Average cell volumes (in μm^3^) were estimated from SSC based on the relationship described by Calvo-Díaz and Morán (2006) assuming spherical shape:

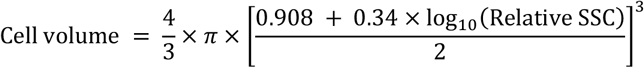

Where *Relative SSC* is the (SSC of prokaryotes)/(SSC of beads) ratio. An in-house calibration was applied to transform relative SSC values from Polysciences-bead-referenced to Molecular-Probes-bead-referenced (Relative SSC_MP_ = 2.201 × Relative SSC_PS_). Prokaryotic biomass was then estimated applying the volumetric relationship (Norland 1993):

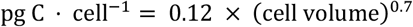

Prokaryotic cell viability was studied *in vivo* right after sample collection by nucleic-acid double-staining (NADS) using SYBR Green I and propidium iodide (Falcioni et al. 2008) following Baltar et al. (2010a). Stained samples were analysed in the cytometer and viable (intact cell membrane) and non-viable (compromised cell membrane) cell groups were identified in FL1 vs FL3 cytograms.

### Prokaryotic activity

Prokaryotic activity was quantified by measuring tritiated leucine incorporation (Kirchman et al. 1985). The centrifugation and filtration methods (Smith and Azam 1992) were applied for samples collected above and below 1000 m depth, respectively. For the centrifugation method, 20 μL of leucine (0.1 μM for 0-150m; 1 μM for 300-1000 m; specific activity of 112 Ci·mmol^−1^) were added to 1 mL of sample (4 replicates, 2 blanks) and these were incubated for 3-12 h. For the filtration method, 200 μL of leucine (1 μM; specific activity of 112 Ci·mmol^−1^) were added to 40 mL of sample (2 replicates, 1 blank) and incubated for 12-24 h. Incubations were stopped by adding trichloroacetic acid to a final concentration of 5% for the centrifugation method and formaldehyde to a final concentration of 2% for the filtration method. For the latter, filters were then washed twice with trichloroacetic acid (50%). After centrifugation/filtration, a scintillation cocktail was added to the samples and the disintegrations per minute (dpm) were computed employing a Wallac scintillation counter with quenching correction, using an external standard. Dpm were converted to leucine incorporation rates based on the equation:

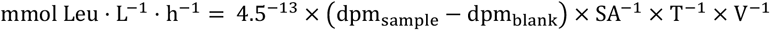

where 4.5^−13^ is the number of curies per dpm (constant), SA is the specific activity of the leucine solution, T is the incubation time in hours and V is the incubation volume in litres. Specific leucine incorporation rates per viable cell were estimated making use of the results from the NADS protocol (see above).

### Multiparameter Water Mass Analysis

The contribution of water masses in each sample deeper than 100 m was objectively quantified using of the optimum multiparameter analysis described in detail in Gómez-Letona et al. (submitted). Briefly, based on previous hydrographic studies in the area (Álvarez et al. 2014; Catalá et al. 2015), twelve source water types were identified in the collected water samples (Fig. S1, Table S1): Salinity Maximum Water (SMW), Madeira Mode Water (MMW), Equatorial Water (EQ_13_), Eastern North Atlantic Central Water of 15ºC (ENACW_15_) and 12ºC (ENACW_12_), Subpolar Mode Water (SPMW), Mediterranean Water (MW), Antarctic Intermediate Water of 5ºC (AAIW_5_) and 3.1ºC (AAIW_3.1_), Circumpolar Deep Water (CDW) and North Atlantic Deep Water of 4.6ºC (NADW_4.6_) and 2ºC (NADW_2_). The contribution of each of them to each sample is quantified by solving a set of four conservative linear mixing equations defined by potential temperature (θ), salinity (S), silicate (SiO_4_H_4_) and the conservative NO tracer (= O_2_ + R_N_·NO_3_ with R_N_ = 9.3; Broecker 1974; Álvarez et al. 2014) (Table S2), plus a fifth equation that constrained the sum of the water mass contributions to 1. See Gómez-Letona et al. (submitted) for more details. The resulting distribution of water masses in the study area can be found in Fig. S1.

Archetype values of physical and biogeochemical variables were estimated for each water mass as weighed means based on the contribution of water masses to each sample (Álvarez-Salgado et al. 2013; Catalá et al. 2015).

### Statistical analyses

All statistical analyses were carried out in R (v. 3.6.0, R Core Team 2019). To assess the relationship between surface productivity and the prokaryotic community, linear regressions were calculated for cytometric variables and leucine incorporation rates using surface Chl-a (average within the upper 20 m) as the independent variable. Regressions were estimated separately for epipelagic (0-200 m), mesopelagic (200-1000 m) and bathypelagic (1000-3000 m) layers, using averaged (for HNA%, cell size, viability, and cell specific leucine incorporation) and integrated (for abundances and bulk leucine incorporation) values of the dependent variables for each depth range. Integrated values were calculated multiplying data values by the distance in meters between data points, based on an interpolated grid (see Fig. 2 and 4) estimated with DIVA (Troupin et al. 2012) in Matlab (R2017a) (see Supporting Methods for details). Regressions of leucine incorporation vs cell size were also calculated.

**Fig. 2.**
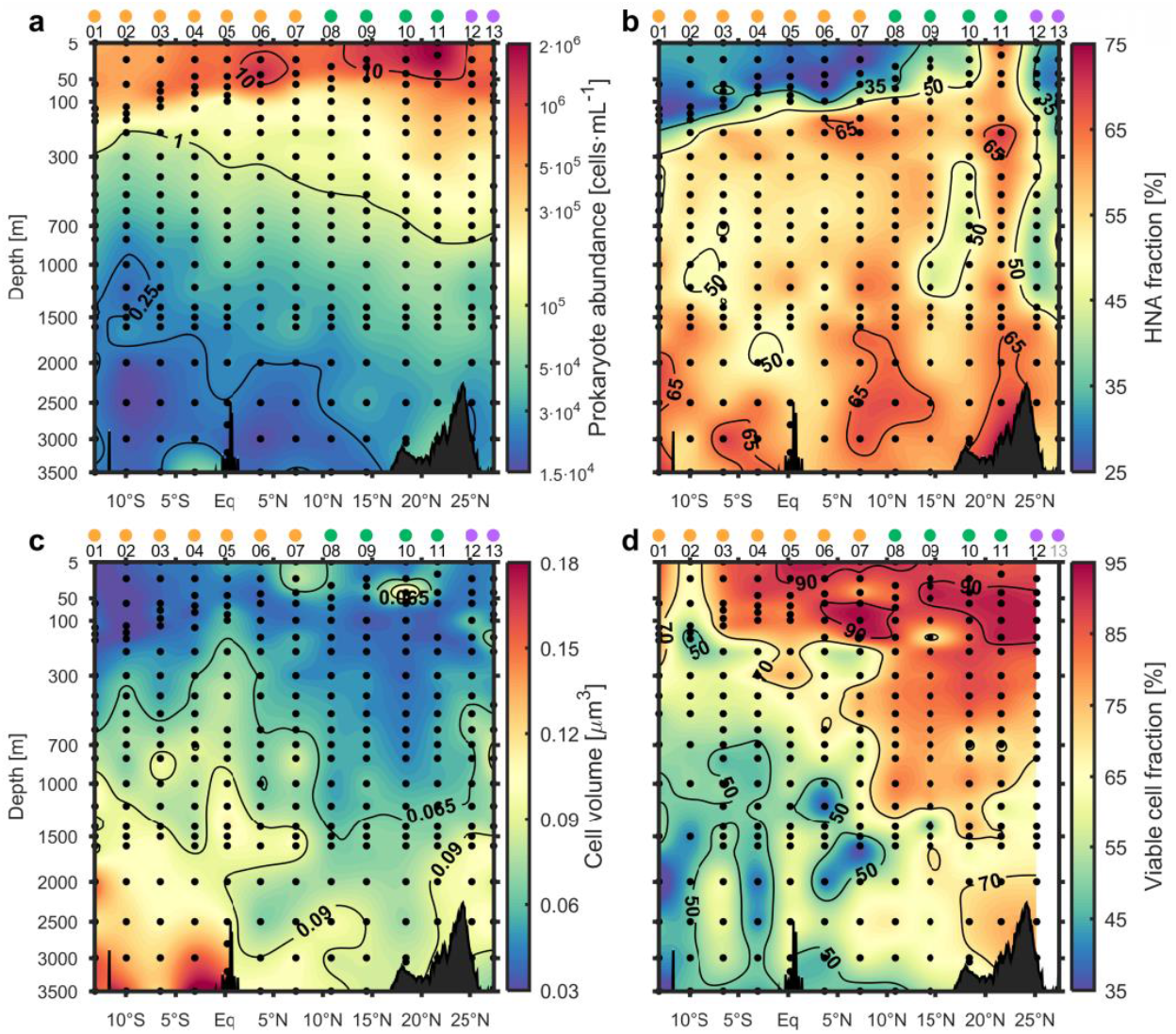
Characterisation of the prokaryotic community by flow cytometry. (a) Total prokaryotic cell abundance. Labels on contour lines are in 10^5^. (b) Fraction of the community represented by HNA prokaryotes. (c) Mean volume of prokaryotic cells. (d) Viability of prokaryotic cells as per NADS (see methods). Black dots represent locations of collected samples and resulting estimates. Dots on top of station numbers represent station groups: ‘South’ (light orange), ‘GD-CB’ (green), ‘North’ (violet). Data interpolation was performed with DIVA in Matlab (R2017a).

Log-log linear regressions of prokaryotic abundances and leucine incorporation vs depth were estimated (including samples of ≥10 m depth) to evaluate vertical trends. Regressions were estimated both jointly for the entire dataset and grouping stations based on surface productivity gradients (see section *Surface productivity gradient along the cruise section* for details): stations 1-7, with low surface Chl-a values, were grouped as ‘South’; stations 8-11, with high Chl-a values, as ‘GD-CB’ (Guinea Dome-Cape Blanc); and stations 12-13, again with low Chl-a values, as ‘North’.

## Results

### Surface productivity gradient along the cruise section

The oceanographic section crossed regions with markedly different productivity regimes, as depicted by net primary production estimates and Chl-a patterns (Fig. 1). Stations in the southern end of the section (1-2), off the coast of Brazil, presented conditions typically associated to oligotrophic waters, with low surface Chl-a values (0.03-0.07 mg·m^−3^) and a Deep Chlorophyll Maximum (DCM) located at ∼135 m depth, showing concentrations that barely exceeded 0.5 mg·m^−3^. Despite consistently showing low surface Chl-a values, stations 3-7 presented an increasing trend, from 0.04-0.06 mg·m^−3^ at station 3 to 0.11-0.19 mg·m^−3^ at station 7. A parallel shoaling and strengthening of the DCM were evident, from 95 m (1.2 mg·m^−3^) at station 3 to 40 m (2.3 mg·m^−3^) at station 7. At stations 8-11 surface Chl-a concentrations markedly increased, transitioning to highly productive waters: from 0.25-1.3 mg·m^−3^ in the Guinea Dome area (stations 8-9) to 1.4-1.5 mg·m^−3^ at station 10 and 4.8-5.9 mg·m^−3^ at Cape Blanc (station 11). These stations showed the shallowest DCMs, with Cape Blanc peaking at 15 m and 5.9 mg·m^−3^. Entering the oligotrophic Canary Current, stations 12-13 presented a sharp decrease in Chl-a values (0.19-0.24 mg·m^−3^), while the DCM became deeper and weaker (85 m and 1.1 mg·m^−3^ at station 13).

### Spatial distribution patterns of cytometric signatures

Prokaryotic abundances (Fig. 2a) showed decreasing numbers of cells with depth in all stations (log-log slope vs depth = -0.816 ± 0.035; Table 1). Concentrations in epipelagic waters were always above 10^5^ cells·mL^−1^ with peaks exceeding 10^6^ cells·mL^−1^ in surface samples in stations 6, 9, 10 and, especially, 11 (Cape Blanc area), where concentrations reached 2·10^6^ cell·mL^−1^. Despite the widespread decrease with depth, prokaryotic abundances displayed latitudinal differences in the dark ocean too: increasing values were observed from the tropical South Atlantic (10^5^ cells·mL^−1^ at ∼300 m and <2.5·10^4^ cells·mL^−1^ below 1000-1500 m) to the Cape Blanc area (10^5^ cells·mL^−1^ at ∼700 m and >2.5·10^4^ cells·mL^−1^ in the entire water column). Indeed, there was a significantly positive relationship between surface Chl-a concentrations and integrated prokaryotic abundances in the epipelagic, mesopelagic, and bathypelagic layers (Fig. 3a, Table S2). These gradients were reflected in changes in log-log slopes (‘South’ = -0.823 ± 0.042 vs ‘GD-CB’ = -0.783 ± 0.045; Table 1; Fig. S2) and the archetype prokaryotic cell abundances of the water masses (Table 2): SMW (134 ± 11 m) in the southern part of the section, presented markedly lower values (27.4 ± 3.2·10^4^ cells·mL^−1^) than MMW (130 ± 17 m; 38.0 ± 5.6·10^4^ cells·mL^−1^), found in the northern part. Similarly, AAIW_5_ (713 ± 47 m) showed lower archetype prokaryotic cell values than SPMW (696 ± 56 m), the former presenting 5.8 ± 0.5·10^4^ cells·mL^−1^ and the latter 9.5 ± 0.8·10^4^ cells·mL^−1^. Bathypelagic water masses were distributed along the entire section and, thus, their differences were only depth-dependent (Table 2), but intra-water mass latitudinal changes were evident (Fig. 2a).

**Fig. 3.**
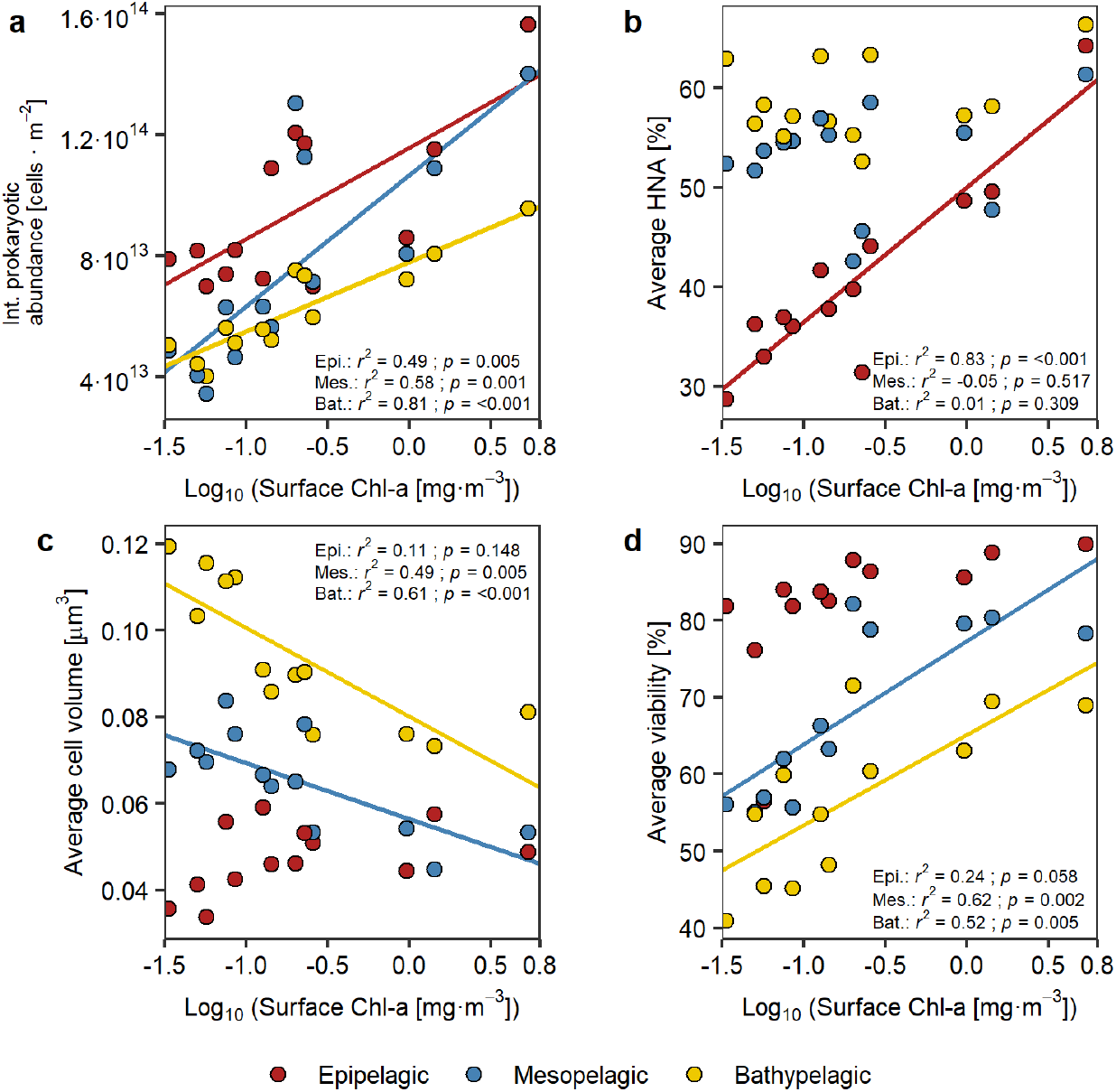
Linear regressions between surface Chl-a concentrations (averaged within the first 20 m, as proxy for productivity) and cytometric variables: (a) integrated prokaryotic abundance, (b) average relative HNA cell abundance, (c) average cell volume, and (d) average cell viability. Regressions estimated separately for epipelagic (≤200 m), mesopelagic (>200 m and ≤1000 m) and bathypelagic (>1000 m and ≤3000 m) layers. Regression lines are only shown for significant (*p* < 0.05) results. Regression parameters are presented in Table S3.

**Table 1.** Log-log regressions of prokaryotic abundances (as cells · mL^−1^) and leucine incorporation rates (as pmol Leu · L^−1^ · d^−1^) vs depth, for samples of ≥10 m. Results are presented for the entire dataset (‘All’) and by station group: ‘South’ (stations 1-7), ‘GD-CB’ (Guinea Dome-Cape Blanc, 8-11) and ‘North’ (12-13, if available). *Slope* and *Intercept* estimates are presented alongside 95% confidence intervals. *r*^*2*^ is the adjusted coefficient of determination, *p* the p-value of the regression and *n* the number of samples per regression.

| Variable | Station group | Slope | Intercept | *r*<sup>2</sup> | *p* | *n* |
| --- | --- | --- | --- | --- | --- | --- |
| HNA prokaryotes | All | -0.691 ± 0.037 | 6.514 ± 0.102 | 0.84 | <0.001 | 261 |
|  | South | -0.661 ± 0.039 | 6.328 ± 0.108 | 0.89 | <0.001 | 142 |
|  | GD-CB | -0.724 ± 0.055 | 6.742 ± 0.151 | 0.90 | <0.001 | 82 |
|  | North | -0.693 ± 0.079 | 6.608 ± 0.215 | 0.90 | <0.001 | 37 |
| LNA prokaryotes | All | -0.923 ± 0.038 | 7.107 ± 0.105 | 0.90 | <0.001 | 261 |
|  | South | -0.953 ± 0.049 | 7.107 ± 0.136 | 0.91 | <0.001 | 142 |
|  | GD-CB | -0.848 ± 0.045 | 6.956 ± 0.125 | 0.95 | <0.001 | 82 |
|  | North | -0.921 ± 0.076 | 7.307 ± 0.207 | 0.94 | <0.001 | 37 |
| All prokaryotes | All | -0.816 ± 0.035 | 7.149 ± 0.096 | 0.89 | <0.001 | 261 |
|  | South | -0.823 ± 0.042 | 7.075 ± 0.117 | 0.91 | <0.001 | 142 |
|  | GD-CB | -0.783 ± 0.045 | 7.153 ± 0.124 | 0.94 | <0.001 | 82 |
|  | North | -0.820 ± 0.069 | 7.305 ± 0.188 | 0.94 | <0.001 | 37 |
| Viable prokaryotes | All | -0.916 ± 0.048 | 7.241 ± 0.131 | 0.86 | <0.001 | 231 |
|  | South | -0.950 ± 0.050 | 7.204 ± 0.140 | 0.92 | <0.001 | 129 |
|  | GD-CB | -0.868 ± 0.058 | 7.255 ± 0.160 | 0.92 | <0.001 | 82 |
|  | North | -0.870 ± 0.118 | 7.354 ± 0.322 | 0.93 | <0.001 | 20 |
| Leucine incorporation | All | -1.261 ± 0.097 | 4.213 ± 0.269 | 0.81 | <0.001 | 152 |
|  | South | -1.331 ± 0.122 | 4.447 ± 0.342 | 0.83 | <0.001 | 98 |
|  | GD-CB | -1.148 ± 0.154 | 3.830 ± 0.424 | 0.81 | <0.001 | 54 |
|  | North | - | - | - | - | - |

**Table 2.** Characterisation of the prokaryotic community in the water masses. Contribution of each water mass to the total sampled volume (as %), and archetype values (± standard error) of depth, latitude and biological parameters.

| Water mass | Volume [%] | Depth [m] | Latitude [°N] | Prokaryotic abundance<sup>a</sup> [cell·mL<sup>-1</sup>] | HNA [%] | Cell viability [%] | Cell volume<sup>b</sup> [$\mu\text{m}^3$] | Prokaryotic biomass [$\mu\text{gC}\cdot\text{L}^{-1}$] | Leucine incorporation [pmol Leu·L<sup>-1</sup>·d<sup>-1</sup>] | Specific leucine incorporation<sup>c</sup> [pmol Leu·viable cell<sup>-1</sup>·d<sup>-1</sup>] |
| --- | --- | --- | --- | --- | --- | --- | --- | --- | --- | --- |
| SMW | 2.88 | 134 ± 11 | -9.68 ± 1.80 | 27.4 ± 3.2 | 31.0 ± 0.0 | 69.4 ± 6.8 | 38.1 ± 4.1 | 3.350 ± 0.386 | 103.93 ± 39.15 | 0.759 ± 0.449 |
| MMW | 1.95 | 130 ± 17 | 22.13 ± 2.67 | 38.0 ± 5.6 | 46.5 ± 0.1 | 89.0 ± 1.7 | 50.4 ± 7.2 | 5.692 ± 1.110 | 27.77 ± 13.02 | 0.092 ± 0.056 |
| EQ<sub>13</sub> | 15.71 | 252 ± 23 | 3.13 ± 1.66 | 15.7 ± 1.4 | 53.7 ± 0.0 | 74.3 ± 2.2 | 50.7 ± 1.9 | 2.195 ± 0.165 | 33.11 ± 11.45 | 0.290 ± 0.119 |
| ENACW<sub>15</sub> | 2.09 | 186 ± 34 | 25.40 ± 1.07 | 35.3 ± 4.8 | 44.3 ± 0.0 | 86.9 ± 1.8 | 54.1 ± 5.8 | 5.527 ± 0.934 | 23.33 ± 34.45 | 0.072 ± 0.166 |
| ENACW<sub>12</sub> | 4.68 | 413 ± 53 | 22.40 ± 1.26 | 21.0 ± 2.4 | 52.8 ± 0.0 | 84.1 ± 2.2 | 57.1 ± 5.0 | 3.186 ± 0.265 | 15.16 ± 7.38 | 0.072 ± 0.043 |
| SPMW | 8.34 | 696 ± 56 | 17.32 ± 1.39 | 9.5 ± 0.8 | 52.4 ± 0.0 | 76.1 ± 2.4 | 58.2 ± 3.4 | 1.493 ± 0.114 | 7.58 ± 1.89 | 0.103 ± 0.029 |
| MW | 0.76 | 1632 ± 386 | 25.28 ± 1.68 | 4.7 ± 1.1 | 54.7 ± 0.1 | 70.8 ± 5.2 | 85.8 ± 12.5 | 0.974 ± 0.257 | 1.65 ± 1.73 | 0.042 ± 0.033 |
| AAIW<sub>5</sub> | 12.83 | 713 ± 47 | 0.22 ± 1.86 | 5.8 ± 0.5 | 53.7 ± 0.0 | 61.4 ± 2.4 | 71.0 ± 3.5 | 1.034 ± 0.086 | 6.34 ± 1.48 | 0.228 ± 0.070 |
| AAIW<sub>3.1</sub> | 0.3 | 1052 ± 315 | -9.26 ± 6.56 | 3.0 ± 0.6 | 55.9 ± 0.1 | 50.1 ± 9.8 | 86.7 ± 25.6 | 0.632 ± 0.201 | 4.01 ± 4.00 | 0.311 ± 0.319 |
| CDW | 5.89 | 1917 ± 232 | 3.92 ± 3.47 | 3.2 ± 0.3 | 58.3 ± 0.0 | 56.1 ± 3.7 | 93.9 ± 6.8 | 0.695 ± 0.072 | 1.99 ± 0.94 | 0.140 ± 0.069 |
| NADW<sub>4.6</sub> | 30 | 1485 ± 75 | 5.99 ± 1.46 | 3.9 ± 0.2 | 56.6 ± 0.0 | 59.3 ± 1.8 | 83.9 ± 2.7 | 0.781 ± 0.038 | 2.45 ± 0.65 | 0.126 ± 0.031 |
| NADW<sub>2</sub> | 14.57 | 2754 ± 98 | 4.66 ± 2.12 | 2.6 ± 0.2 | 61.7 ± 0.0 | 55.7 ± 2.3 | 107.8 ± 5.4 | 0.646 ± 0.058 | 1.08 ± 0.13 | 0.086 ± 0.013 |

<sup>a</sup> $\times 10^4$
<sup>b</sup> $\times 10^{-3}$
<sup>c</sup> $\times 10^{-6}$

Prokaryotic biomass (Fig. S3) was largely governed by cell abundances. Surface values increased from 5-7 *μ*gC·L^−1^ in the south to 15-20 *μ*gC·L^−1^ towards the Guinea Dome area, peaking at Cape Blanc at 36 *μ*gC·L^−1^. As abundance patterns, while decreasing with depth, biomass also displayed a latitudinal gradient in both the mesopelagic and bathypelagic layers, producing significant positive relationships with Chl-a in epi-, meso- and bathypelagic layers (Table S3). Resulting archetype values for AAIW_5_ and SPMW were 1.03 ± 0.09 and 1.49 ± 0.11 *μ*gC·L^−1^, respectively, while in bathypelagic waters biomasses below 0.55 *μ*gC·L^−1^ in the south contrasted with values of 0.6-1 *μ*gC·L^−1^ in the north.

Surface communities were dominated by LNA cells (in many cases exceeding 65% of counts) except in station 11 (Fig. 2b). An increase of HNA contribution was observed from the southern stations across the Guinea Dome area towards Cape Blanc. In the dark ocean, HNA prokaryotes overall dominated the community (Fig. 2b), their contribution increasing with depth and reaching >65% in some bathypelagic samples. This was evident too from the different log-log slopes (Table 1), as the abundance of LNA prokaryotes (-0.923 ± 0.038) decreased markedly faster than HNA prokaryotes (-0.691 ± 0.037). While HNA contribution in station 11 was >50% in the entire water column (with particularly high values (75%) at the bottom), in stations 12 and 13 HNA contributions were back below 50% (down to ∼1500 m). However, no significant relationship was observed between surface Chl-a and average HNA % in the dark ocean, only the epipelagic layer showing a significant positive relationship (Fig. 3b, Table S3).

The average cell volume (Fig. 2c) of prokaryotic communities was lowest in surface waters, with values of 0.030-0.060 *μ*m^3^ that were rather uniform along the entire cruise section, resulting in no significant relationship with Chl-a concentrations (Fig. 3c, Table S3). Cell volumes overall increased with depth, although with clear latitudinal differences: in stations 1-7 and 12-13, volumes of >0.065 *μ*m^3^ were reached between 300-700 m depth, while in stations 8-11 such volumes were only present below 1200-1500 m. These latitudinal differences in deep waters were reflected in significant negative relationships with surface Chl-a concentrations, both in the meso- and bathypelagic layers (Fig. 3c, Table S3). This was also evident in archetype cell volumes of water masses, e.g., AAIW_5_ and SPMW presented values of 0.0710 ± 0.0035 and 0.0582 ± 0.0034 *μ*m^3^, respectively (Table 2). The highest average cell volumes were observed in bathypelagic waters of stations 1-7, most samples exceeding 0.090 *μ*m^3^ (some reaching >0.15 *μ*m^3^)

The fraction of viable cells (presenting intact cell membranes) was highest in epipelagic waters with widespread >70% contributions to the total number of detected cells and peaks exceeding 90% (Fig. 2d). Viability however consistently decreased with depth along the section (log-log slopes of -0.916 ± 0.048, Table 1), following the opposite pattern to cell volume. Stations 1-7 showed quick decreases in the abundance of viable cells (log-log slope = -0.950 ± 0.050; Table 1, Fig. S2), with proportions <70% immediately below the epipelagic layer, and <50% below ∼1000 m in many samples, especially in stations 1-4. In stations 8-12 on the contrary (log-log slope in GD-CB = -0.868 ± 0.058; Table 1, Fig. S2), viabilities above 70% were measured down to ∼1000 m. This produced archetype viabilities of 61.4 ± 2.4 % and 76.1 ± 2.4 for AAIW_5_ and SPMW, respectively. This gradient along the section was consistent down to 3500 m, were values of 35-50% in stations 1-2 contrasted with >70% in samples close to the bottom at stations 10-12. These changes yielded significant positive relationships between surface Chl-a and cell viability in the meso- and bathypelagic waters (Fig. 3d, Table S3). Epipelagic viability also displayed a positive relationship with surface Chl-a (Fig. 3d), although it was not significant.

Leucine incorporation rates (Fig. 4a) decreased drastically with depth (log-log slope = - 1.261 ± 0.097; Table 1). Overall leucine incorporation rates were greater than 150 pmol Leu·L^−1^·d^−1^ in the upper 50 m of the water column, with peaks at stations 4 (564 pmol Leu·L^−1^·d^−1^), 7 (420 pmol Leu·L^−1^·d^−1^), 9 (377 pmol Leu·L^−1^·d^−1^), and 11 (823 pmol Leu·L^−1^·d^−1^). Rates decreased to 15 pmol Leu·L^−1^·d^−1^ by 300 m and to 2 pmol Leu·L^−1^·d^−1^ by 1000-1500 m depth. No consistent latitudinal differences were observed in mesopelagic samples, but overall leucine incorporation rates in the bathypelagic were higher in the south, with the exception of samples close to the bottom at stations 9-11. Log-log slopes vs. depth in ‘South’ and ‘GD-CB’ stations, although different, had considerable confidence intervals (-1.331 ± 0.122 and -1.148 ± 0.154, respectively; Table 1), and no significant relationships were found between leucine incorporation rates and surface Chl-a concentrations (Fig. S4a, Table S3).

Specific leucine incorporation rates per viable cell (Fig. 4b) tended to be higher in the southern end of the section. In epipelagic waters specific rates were widely above 0.3·10^−6^ pmol Leu·viable cell^−1^·d^−1^ in stations 1-5, with multiple samples exceeding 0.6 pmol Leu·viable cell^−1^·d^−1^. In stations 8-11 values were lower and barely exceed 0.3·10^−6^ pmol Leu·viable cell^−1^·d^−1^. In mesopelagic samples specific leucine incorporation rates per viable cell ranged between (0.06-0.3)·10^−6^ pmol Leu·viable cell^−1^·d^−1^ and were slightly greater in the south, although no clear differences were observed between water masses (SPMW = (0.103 ± 0.029)·10^−6^ and AAIW_5_ = (0.228 ± 0.070)·10^−6^ pmol Leu·viable cell^−1^·d^−1^, Table 2). As bulk leucine incorporation rates, viable cell specific rates in bathypelagic waters tended to increase towards the south, overall exceeding 0.06·10^−6^ pmol Leu·viable cell^−1^·d^−1^, while stations 8-11 mostly showed specific rates below 0.06·10^−6^ pmol Leu·viable cell^−1^·d^−1^, except samples close to the bottom. Average viable cell specific rates in bathypelagic waters did show a (weak) negative relationship with surface Chl-a (Fig. S4b, Table S3).

**Fig. 4.**
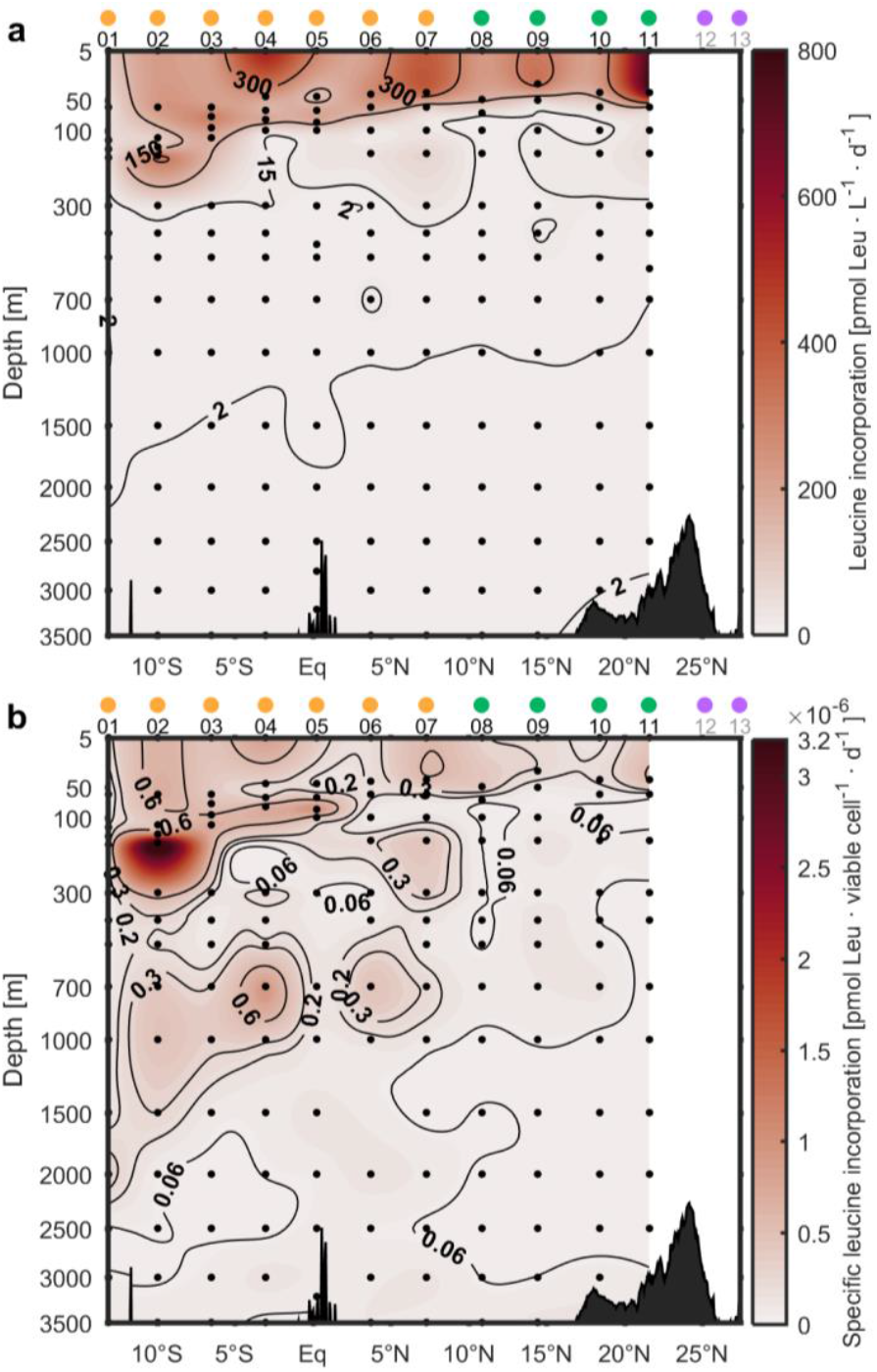
Prokaryotic heterotrophic production. (a) Bulk leucine incorporation rate, (b) Specific leucine incorporation rate per viable cell (contour lines are in 10^−6^). Black dots represent locations of collected samples and resulting estimates. Dots on top of station numbers represent station groups: ‘South’ (light orange), ‘GD-CB’ (green), ‘North’ (violet). Data interpolation was performed with DIVA in Matlab (R2017a).

## Discussion

The oceanographic section studied encompassed widely diverse environments along the tropical and subtropical Atlantic, from the oligotrophic tropical south Atlantic to the highly productive waters of the Cape Blanc area in the Northwest Africa upwelling system (Fig. 1a; Carr and Kearns 2003). This transition from waters of low to high productivity was well captured by the increasing concentrations of Chl-a measured in epipelagic waters from stations 1 to 11, along with a shoaling of the DCM (Fig. 1b). Based on results obtained during the same cruise, Gómez-Letona et al. (submitted) reported intense fluorescence signals of labile, protein-like dissolved organic matter at stations 8, 9, 11 and 12 for the entire water column. These findings suggest a remarkable vertical transport of organic matter from the surface into the dark ocean in the Guinea Dome-Cape Blanc region, in agreement with previous studies with sediment traps reporting high sinking particle fluxes (Fischer et al. 2020).

The vertical decrease in cell abundances of the different prokaryotic groups (total, LNA, HNA, and viable cells), evaluated with log-log relationship vs. depth, yielded slope values that were in the range of those reported in the literature (Gasol et al. 2009, 2019; Arístegui et al. 2009). Trends of vertical decreases in total prokaryotic cell abundances and the relative abundance of viable cells are widespread in the ocean (Gasol et al. 2009, 2019) and point to harsher conditions (low temperature, high pressure, reduced DOM availability) for prokaryotes in deep ocean environments (Herndl and Reinthaler 2013). Increases in the proportion of HNA prokaryotes and cell volume with depth have also been reported previously (Van Wambeke et al. 2011), and can be interpreted as an increase in the contribution of larger prokaryotic cells, with bigger genomes (Bouvier et al. 2007). Besides the vertical variability, latitudinal gradients were observed in the deep ocean, matching surface productivity gradients, as defined by Chl-a concentrations. The tendency to more abundant, more viable and smaller cells in the mesopelagic and bathypelagic layers under highly productive surface waters with potentially important vertical flux of particles (Fischer et al. 2020) suggests a remarkable effect of sinking particles in the prokaryotic communities of the dark ocean. In particular, such relationship between surface productivity and prokaryotic viability in deep waters has not been previously reported to the authors’ knowledge. The effect of vertical connectivity would be twofold, including the transport of both resources and microorganisms. Particles would potentially introduce fresh organic matter, as suggested by the vertical link displayed by protein-like (resembling peak T, Coble 1996) fluorescent dissolved organic matter and surface productivity proxies in the study area (Gómez-Letona et al., submitted), like in other oceanic regions (Ruiz-González et al. 2020), fuelling prokaryotes and allowing them to attain higher viabilities and abundances (Hansell and Ducklow 2003; Yokokawa et al. 2013). Moreover, the sinking particles could also act as vectors, transporting particle-attached prokaryotes from epipelagic communities to the dark ocean, yielding communities more similar to each other than in regions with low vertical transport (Mestre et al. 2018). This could explain the smaller size of deep ocean prokaryotes under the more productive stations.

The log-log slope of leucine incorporation rates for the entire dataset was in the range of others reported in the literature (Arístegui et al. 2009). An exception to the decreasing rates of leucine incorporation with depth was found in samples close to the bottom at stations 9-11, which showed relatively high rates that might have been fuelled by resuspension (Fischer et al. 2009; Ziervogel et al. 2016). We found absent (for bulk rates) or negative (for cell specific rates in the bathypelagic) relationships between surface productivity and leucine incorporation in the dark ocean (Fig. S4, Table S3), manifesting that, while the vertical connection between surface productivity and prokaryotic abundance was evident, the connection with heterotrophic production is unclear. These findings are in line with previous studies that found positive but weak relationships between vertical particle flux and prokaryotic production in the mesopelagic, but none in the bathypelagic (Yokokawa et al. 2013).

These unclear relationships may be due to methodological limitations, since prokaryotic production is estimated from metabolic rates, but sometimes metabolism is not directed to cell-growth but to cell maintenance (Carlson et al. 2007; Giering and Evans 2022). Particularly in hostile environmental conditions, with low resources and energy supply, prokaryotes have been found to achieve lower growth efficiencies (del Giorgio and Cole 1998). In these situations, resources and energy are directed in greater proportions to the maintenance of cell structures and functionality, instead of biomass production and growth (Carlson et al. 2007). The bathypelagic waters of stations 1-4 showed minimum cell viability ratios, in agreement with expected harsher conditions due to potential low external inputs of organic substrates in these oligotrophic waters. The relatively high specific leucine incorporation rates in the deep waters of these southern stations might thus be linked to metabolism directed to maintenance, rather than growth and cell division. Larger prokaryotes have greater metabolic requirements than smaller ones (del Giorgio and Gasol 2008), and we indeed found that high average values of specific leucine incorporation in the bathypelagic were associated with high average cell sizes (Figs. 2, 3 and S5). In fact, there is increasing evidence that leucine-to-carbon conversion factors are highly variable in the ocean, decreasing with depth and increasing with water productivity (del Giorgio and Cole 1998; Orta-Ponce et al. 2021; Giering and Evans 2022). Considering these trends, the negative relationship between leucine incorporation in bathypelagic samples and surface productivity would not directly translate to prokaryotic heterotrophic production, as lower conversion factors could be expected in the southern stations relative to the Guinea Dome – Cape Blanc area. This would potentially result in higher production rates under the productive waters, which would agree with the other trends of increased viability and cell numbers. Thus, variation in the leucine-to-carbon conversion factors probably explain the lack of relationship between surface productivity and prokaryotic heterotrophic activity (as measured by leucine incorporation) in the dark realm.

## Conclusions

We studied the abundance, cytometric signatures and heterotrophic metabolism of prokaryotic communities from surface down to 3500 m depth along a primary productivity gradient in the subtropical and tropical Atlantic. Our results show that deeper waters tended to harbour communities with reduced cell concentrations and viability, but larger cell sizes and genomes. The trends were coupled with changes along surface productivity: waters under highly productive areas presented higher cell counts, biomass and viability, and lower average cell sizes, than waters under oligotrophic zones. These relationships were significant down to the bathypelagic zone (and were reflected in the archetype values of water masses), highlighting the extent of the vertical connectivity along the water column. The heterotrophic metabolism, however, was not equally affected: while bulk leucine incorporation rates displayed no significant relationship with surface productivity, cell specific rates showed a negative relationship in bathypelagic waters. Energy and resource allocation to cellular maintenance processes under adverse environmental conditions (resulting in low conversion of the incorporated leucine into biomass), and dependence of metabolic requirements on cell size could influence the conversion of leucine incorporation into carbon production. Our work underlines the strong link between the productivity of surface layers and the biomass and physiological status of deep ocean prokaryotic communities.

## Acknowledgements

We thank to the officers and crew of the *BIO Hespérides*, and the staff of the Unit of Marine Technology (UTM) of the Spanish Research Council (CSIC) for their invaluable help at sea.

This work is a contribution to projects MAFIA (grant number CTM2012-39587-C04-01), FLUXES (grant number CTM2015-69392-C3), e-IMPACT (grant number PID2019-109084RB-C21), and INTERES (CTM2017-83362-R) funded by the Spanish “Plan Nacional/Estatal de I+D” and cofounded with FEDER funds, and to project SUMMER (grant number AMD-817806-5) funded by the European Union’s Horizon 2020 research and innovation program. MGL is supported by Ministerio de Ciencia, Innovación y Universidades, Gobierno de España (grant number FPU17-01435) during his PhD. MS is supported by the Project MIAU (grant number RTI2018-101025-B-I00) and the ‘Severo Ochoa Centre of Excellence’ accreditation (CEX2019-000928-S).

