## Supporting material for "Surface productivity gradients govern changes in abundance and physiological status of deep ocean prokaryotes across the tropical and subtropical Atlantic"

This file contains:

Supporting Figures 1-5

Supporting Tables 1-3

Supporting Methods

References

### Supporting figures

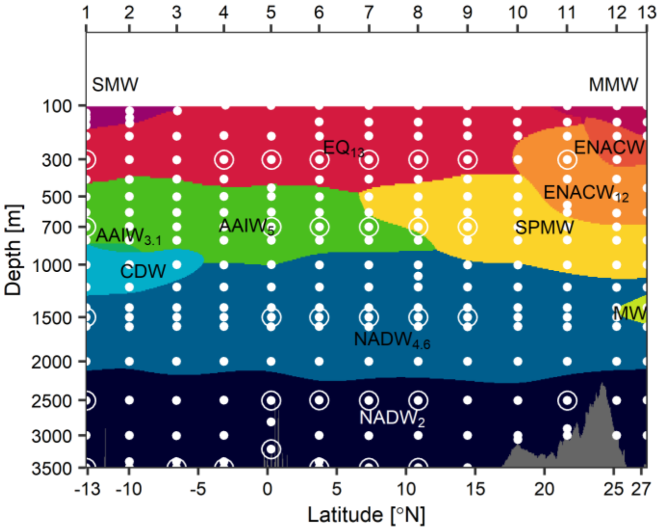


Fig. S1. Distribution of the water masses in the study section. Acronyms are as follows: Salinity Maximum Water (SMW), Madeira Mode Water (MMW), Equatorial Water ($EQ_{13}$), Eastern North Atlantic Central Water of 15ºC ($ENACW_{15}$) and 12ºC ($ENACW_{12}$), Subpolar Mode Water (SPMW), Mediterranean Water (MW), Antarctic Intermediate Water of 5ºC ($AAIW_5$) and 3.1ºC ($AAIW_{3.1}$), Circumpolar Deep Water (CDW) and North Atlantic Deep Water of 4.6ºC ($NADW_{4.6}$) and 2ºC ($NADW_2$). Approximate area in which each water mass has their highest contribution is shown. Dots represent samples of the physical and geochemical variables included in the optimum multiparameter analysis. Note that the vertical scale is square root-transformed to allow for a better visualisation of the results. Numbers on top correspond to stations in Fig. 1 in the main text. Data interpolation was performed with DIVA in Matlab (R2017a). Modified from Gómez-Letona et al. (submitted).

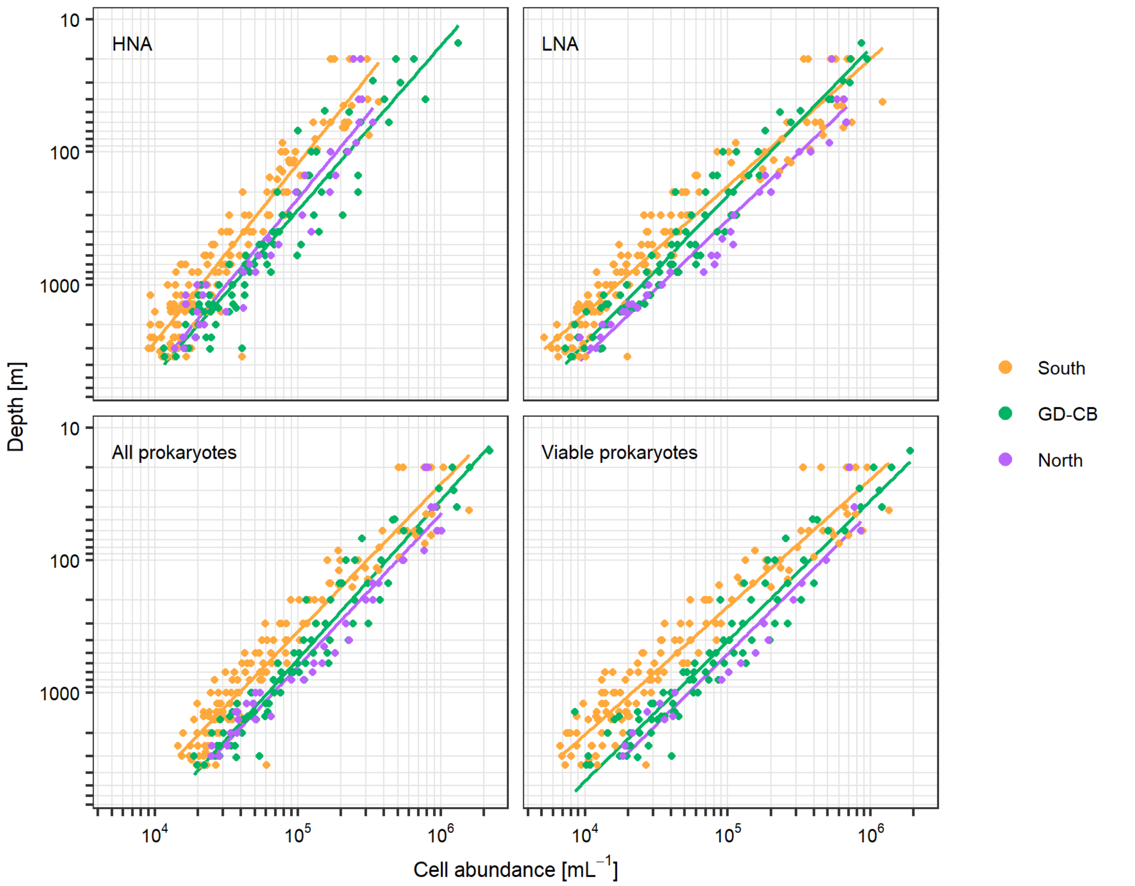


Fig. S2. Log-log regressions of prokaryotic abundances vs depth (for samples of ≥10 m), by station group: 'South' (stations 1-7; light orange), 'GD-CB' (Guinea Dome-Cape Blanc, 8-11; green) and 'North' (12-13, if available; violet). The regression parameters can be found in Table 1 in the main text.

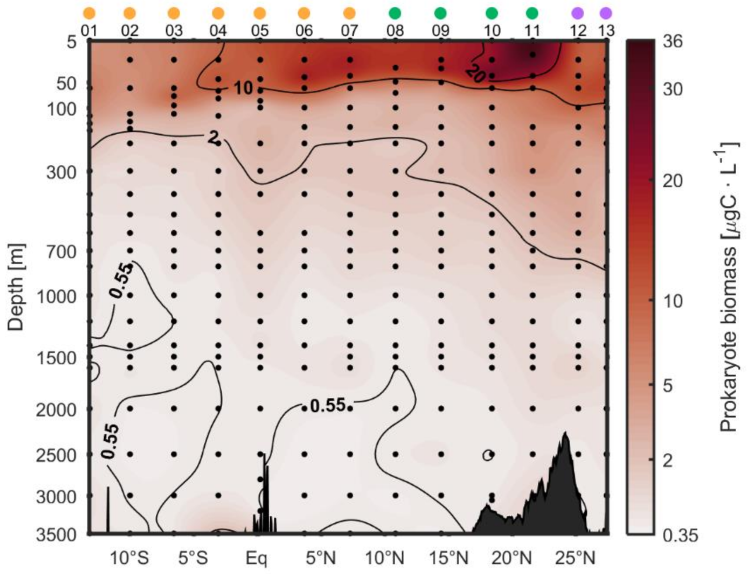


Fig. S3. Biomass of prokaryotes. Black dots represent collected samples. Dots on top of station numbers represent station groups: ‘South’ (light orange), ‘GD-CB’ (green), ‘North’ (violet). Data interpolation was performed with DIVA in Matlab (R2017a).

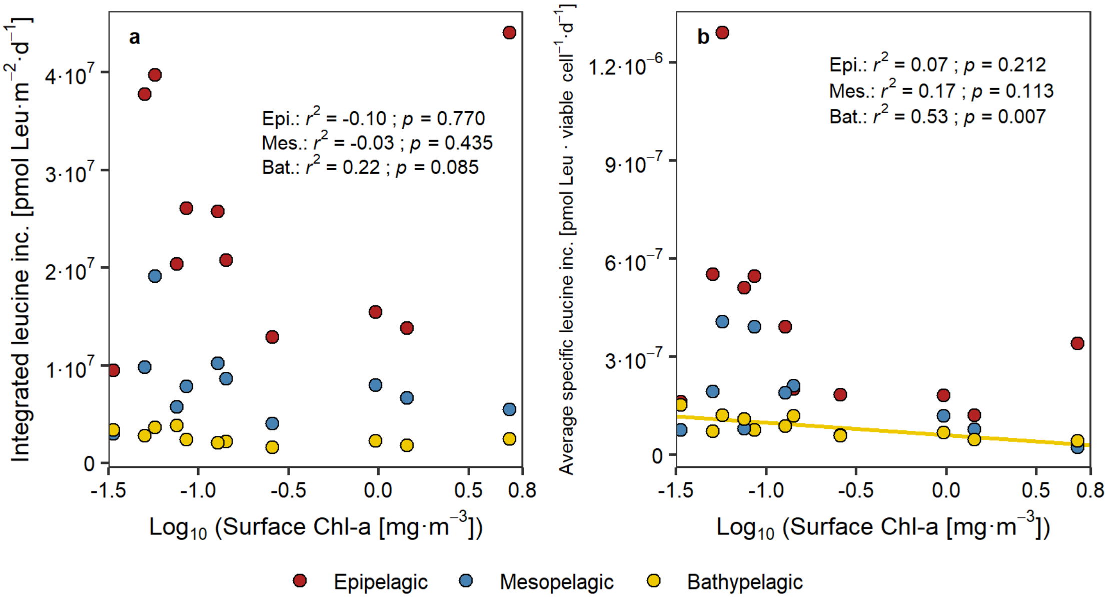


Fig. S4. Linear regressions between surface Chl-a concentrations (averaged within the first 20 m, as proxy for productivity) and metabolic variables: A) integrated leucine incorporation rate, B) average viable cell specific leucine incorporation rate. Regressions estimated separately for epipelagic (≤200 m), mesopelagic (>200 m and ≤1000 m) and bathypelagic (>1000 m and ≤3000 m) layers. Regression lines are only shown for significant ($p$ < 0.05) results. Regression parameters are presented in Table S3.

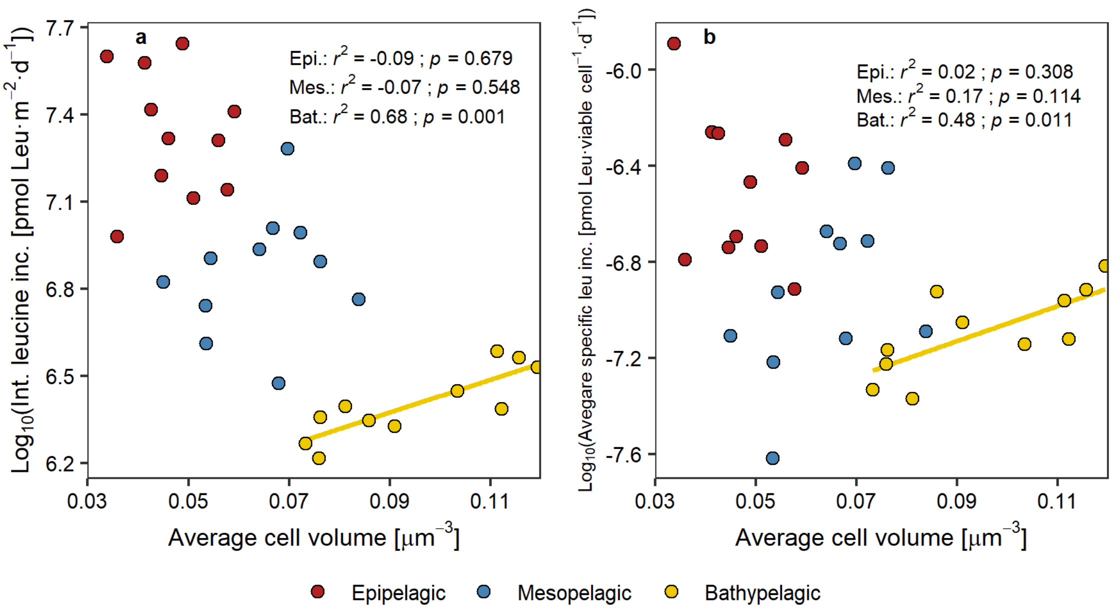


Fig. S5. Linear regression between surface average cell volume and A) integrated leucine incorporation rate, and B) average viable cell specific leucine incorporation, estimated separately for epipelagic (≤200 m), mesopelagic (>200 m and ≤1000 m) and bathypelagic (>1000 m and ≤3000 m) layers. Regression lines are only shown for significant ($p < 0.05$) results.

### Supporting tables

**Supporting Table 1.** Water types intercepted during the MAFIA cruise, brief description of source point where they belong to, characteristics, and some references with more details about their origin and circulation.

| Name and acronym | Source | Characteristics | SWT | References |
|---|---|---|---|---|
| Salinity Maximum Water (SMW) | Tropical area (12–22ºS) | Warmest (27ºC) mode water in the SAO, formed by evaporation, transported westward to America with SEC | SMW | Worthington (1976), Stramma and England (1999), Mémery et al. (2000) |
| Equatorial Atlantic Central Water (13ºC) | Eastern South Atlantic, near Namibia | Formed by mixing of low salinity water outcropped further south with overlying high salinity water. Transported by the South Equatorial current to the Equator and along the Brazilian coast by the North Brazil current. | $EQ_{13}$ | Tsuchiya (1986) |
| MMW | Madeira Mode Water | Mode water formed near the Madeira Island | MMW | Siedler et al. (1987) |
| Eastern North Atlantic Central water | Eastern North Atlantic Subtropical gyre | Mode waters defining the upper (15ºC) and lower (12ºC) limits of the subtropical ENACW formed between the area of the Azores and Portugal currents. | $ENACW_{15}$ $ENACW_{12}$ | Harvey (1982), Pollard and Pu (1985), Ríos et al. (1992), Álvarez-Salgado et al. (2013) |
| Mediterranean Water | Gulf of Cadiz | Formed in the Gulf of Cadiz by entrainment of Eastern North Atlantic Central water on the high-salinity outflow from the Mediterranean Sea, spreads at 800–1300 m, S > 36 and θ ~ 11–12 ºC. | MW | Zenk (1975), Ambar and Howe (1979), Castro et al. (1998), Álvarez-Salgado et al. (2013) |
| Antarctic Intermediate Water | Pacific Ocean north of the Sub-Antarctic Front & Malvinas-Brazil Confluence. | Formed north of the Subantarctic Front (SAF) and east of the Drake Passage by ventilation of the Subantarctic Mode Water (SAMW) formed in the Southeast Pacific. | $AAIW_{5.0}$ $AAIW_{3.1}$ | McCartney (1982), Piola and Gordon (1989), Talley (1996) |
| Circumpolar Deep Water | Antarctic Circumpolar Current | Also named Common Water, formed by mixing in the Antarctic Circumpolar current of mid-depth Indian, Pacific and Atlantic deep water with WSDW and NADW. | CDW | Montgomery (1958), Georgi (1981), Broecker et al. (1985) |
| North Atlantic Deep Water | North Atlantic Ocean | Carried into the South Atlantic by the Deep Western Boundary Current (DWBC). Characterized by salinity maximum and silicate minimum (4.6ºC); and θ–S discontinuity and oxygen maximum (2.0 ºC). Defined at their entry in the South Atlantic Ocean off South America. | $NADW_{4.6}$ $NADW_{2.0}$ | Wüst (1935), Speer and McCartney (1992), Friedrichs et al. (1994) |

**Supporting Table 2.** Thermohaline and chemical characteristics (average value ± uncertainty) of the water types (WT) introduced in the OMP analysis of the water masses intercepted during the MAFIA cruise.

| WT | $\theta_i$ (ºC) | $S_i$ | $SiO_4H_{4i}$ (µmol kg$^{-1}$) | $NO_i$ (µmol kg$^{-1}$) |
|---|---|---|---|---|
| SMW [a] | 27.0 ± 0.1 | 37.50 ± 0.01 | 1.1 ± 0.5 | 206 ± 3 |
| MMW [b] | 20.0 ± 0.5 | 37.00 ± 0.04 | 0.4 ± 0.3 | 225 ± 10 |
| $EQ_{13}$ [a] | 13.0 ± 0.1 | 35.20 ± 0.01 | 5.3 ± 0.7 | 315 ± 3 |
| $ENACW_{15}$ [b] | 15.3 ± 0.4 | 36.10 ± 0.02 | 2.2 ± 1.7 | 264 ± 8 |
| $ENACW_{12}$ [c] | 12.2 ± 0.4 | 35.66 ± 0.02 | 4.9 ± 0.2 | 322 ± 8 |
| SPMW [c] | 8.2 ± 0.4 | 35.23 ± 0.01 | 14.5 ± 0.4 | 386 ± 7 |
| MW [c] | 11.8 ± 0.1 | 36.50 ± 0.01 | 7.2 ± 0.7 | 304 ± 9 |
| $AAIW_5$ [a] | 5.00 ± 0.08 | 34.14 ± 0.01 | 7.0 ± 0.7 | 482 ± 3 |
| $AAIW_{3.1}$ [a] | 3.10 ± 0.08 | 34.12 ± 0.01 | 16.4 ± 0.7 | 558 ± 3 |
| CDW [a] | 1.60 ± 0.03 | 34.720 ± 0.003 | 110.6 ± 0.9 | 497 ± 1 |
| $NADW_{4.6}$ [a] | 4.6 ± 0.1 | 35.020 ± 0.005 | 7.3 ± 0.5 | 426 ± 2 |
| $NADW_2$ [a] | 2.02 ± 0.03 | 34.910 ± 0.003 | 28.2 ± 0.9 | 446 ± 1 |

[a] Álvarez et al. (2014)
[b] Álvarez and Álvarez-Salgado (2009); Lønborg and Álvarez-Salgado (2014)
[c] Pérez et al. (2001); Álvarez and Álvarez-Salgado (2009)

**Supporting Table 3**. Results from the regressions of cytometric and metabolic variables vs log-transformed surface Chl-a values. Regressions were performed by layer, with integrated values for variables with units referenced to volume and averaged values otherwise. *Slope* and *Intercept* estimates are presented alongside 95% confidence intervals. $r^2$ is the adjusted coefficient of determination, *p* the p-value of the regression. Regressions are shown in Figs. S4 and S6.

| | | Layer | Slope | Intercept | $r^2$ | *p* |
|---|---|---|---|---|---|---|
| | Prokaryotic abundance | Epipelagic | $(3.02 \pm 1.89)\cdot10^{13}$ | $(1.16 \pm 0.18)\cdot10^{14}$ | 0.49 | 0.005 |
| | | Mesopelagic | $(4.34 \pm 2.27)\cdot10^{13}$ | $(1.07 \pm 0.21)\cdot10^{14}$ | 0.58 | 0.001 |
| | | Bathypelagic | $(2.29 \pm 0.69)\cdot10^{13}$ | $(7.81 \pm 0.64)\cdot10^{13}$ | 0.81 | <0.001 |
| | HNA% | Epipelagic | 13.56 ± 3.86 | 50.06 ± 3.58 | 0.83 | <0.001 |
| | | Mesopelagic | 1.62 ± 5.31 | 54.25 ± 4.91 | -0.05 | 0.517 |
| | | Bathypelagic | 1.93 ± 3.98 | 60.04 ± 3.69 | 0.01 | 0.309 |
| | Cell volume | Epipelagic | 0.0053 ± 0.0075 | 0.051 ± 0.007 | 0.11 | 0.148 |
| | | Mesopelagic | -0.013 ± 0.008 | 0.056 ± 0.007 | 0.49 | 0.005 |
| | | Bathypelagic | -0.020 ± 0.010 | 0.080 ± 0.009 | 0.61 | <0.001 |
| | Viability | Epipelagic | 7.47 ± 7.79 | 87.35 ± 7.36 | 0.24 | 0.058 |
| $Log_{10}(Chl\text{-}a_{srf})$ | | Mesopelagic | 13.43 ± 6.94 | 77.31 ± 6.56 | 0.62 | 0.002 |
| | | Bathypelagic | 11.74 ± 7.25 | 65.13 ± 6.85 | 0.52 | 0.005 |
| | Biomass | Epipelagic | (5.30 ± 2.72)·105 | (1.70 ± 0.25)·106 | 0.59 | 0.001 |
| | | Mesopelagic | (5.41 ± 4.30)·105 | (1.64 ± 0.40)·106 | 0.36 | 0.018 |
| | | Bathypelagic | (3.08 ± 2.14)·105 | (1.57 ± 0.20)·106 | 0.43 | 0.009 |
| | Leucine incorporation rate | Epipelagic | $(1.68 \pm 12.64)\cdot10^{6}$ | $(2.54 \pm 1.22)\cdot10^{7}$ | -0.10 | 0.770 |
| | | Mesopelagic | $(-1.63 \pm 4.51)\cdot10^{6}$ | $(6.93 \pm 4.35)\cdot10^{6}$ | -0.03 | 0.435 |
| | | Bathypelagic | $(-5.66 \pm 6.62)\cdot10^{5}$ | $(2.22 \pm 0.64)\cdot10^{6}$ | 0.22 | 0.085 |
| | Leucine incorporation rate per viable cell | Epipelagic | $(-1.96 \pm 3.30)\cdot10^{-7}$ | $(2.71 \pm 3.19)\cdot10^{-7}$ | 0.07 | 0.212 |
| | | Mesopelagic | $(-9.41 \pm 12.13)\cdot10^{-8}$ | $(1.01 \pm 1.17)\cdot10^{-7}$ | 0.17 | 0.113 |
| | | Bathypelagic | $(-3.81 \pm 2.45)\cdot10^{-8}$ | $(6.06 \pm 2.36)\cdot10^{-8}$ | 0.53 | 0.007 |

### Supporting methods

*Inorganic nutrients*

Inorganic nutrients (included in the OMP analysis) were sampled from Niskin bottles with polyethylene tubes and stored at −20°C until analysis in the laboratory. The analysis was performed with a QuAAtro 39-SEAL Analytical AutoAnalyzer following Armstrong et al. (1967).

*Data interpolation*

Interpolations of discrete data were performed with DIVA (Troupin et al. 2012) in Matlab (R2017a). As samples were distributed along different depth scales in epipelagic/upper-mesopelagic waters relative to meso-/bathypelagic waters, the interpolation was done separately for depths 5-400 m ("upper layer") and 300-3500 m ("lower layer"), allowing to fine tune the interpolation appropriately for each depth scale. For the interpolation of the upper layer, the interpolation grid was [5:1:100 105:5:200 210:10:400], with horizontal (LX) and vertical (LY) *Length scales* being 4 and 50, respectively, and the *signal to noise ratio* (SN) 16. For the interpolation of the lower layer the grid was [300:10:1000 1050:50:3500], LX = 4, LY = 500 and SN = 16. Both grids were combined into a single one by a weighted mean of interpolated values in the common depth range (300-400 m): for 300 m, the upper grid was weighted 1 and the lower 0; for 350 m they were equally weighted 0.5; for 400 m 0 and 1, respectively; etc.
